# Focused screening reveals functional effects of microRNAs differentially expressed in colorectal cancer

**DOI:** 10.1101/601484

**Authors:** Danuta Sastre, João Baiochi, Ildercilio Mota de Souza Lima, Josiane Lilian dos Santos Schiavinato, Dimas Tadeu Covas, Rodrigo Alexandre Panepucci

## Abstract

**Background:** Colorectal cancer (CRC) is still a leading cause of death worldwide. Recent studies have pointed to an important role of microRNAs carcinogenesis. In fact, several microRNAs have been described as aberrantly expressed in CRC tissues and in the serum of patients. More specifically, microRNAs with dual roles in both cancer and stem cell survival represent a potential source of novel molecular targets in CRC due to their described functions in normal and deregulated proliferation. However, the functional outcomes of microRNA aberrant expression still need to be explored at the cellular level. Here, we aimed to investigate the effects of microRNAs involved in the control of pluripotency of stem cells in the proliferation and cell death of a colorectal cancer cell line.

**Methods:** We performed transfection of 31 microRNA mimics in HCT116 CRC cells. Cell proliferation and cell death were measured after 4 days of treatment using fluorescence staining in a high content screening platform. Total number of live and dead cells were automatically counted and analyzed. To reveal mRNA targets, we used an oligonucleotide microarray. Functional classification of targets was done using DAVID tool. Gene expression of potential mRNA targets was performed by qPCR.

**Results:** Twenty microRNAs altered the proliferation of HCT116 cells in comparison to control. Three microRNAs significantly repressed cell proliferation and induced cell death simultaneously (miR-22-3p, miR-24-3p, and miR-101-3p). Interestingly, all anti-proliferative microRNAs in our study had been previously described as poorly expressed in the CRC samples and were implicated in the disease. Microarray analysis of miR-101-3p targets revealed Wnt and cancer as pathways regulated by this microRNA. Specific repression of anti-apoptotic isoform of MCL-1, a member of the BCL-2 family, was also identified as a possible mechanism for miR-101-3p anti-proliferative/pro-apoptotic effect.

**Conclusions:** microRNAs described as upregulated in CRC tend to induce proliferation in vitro, whereas microRNAs described as poorly expressed in CRC halt proliferation and induce cell death in vitro. Selective inhibition of anti-apoptotic MCL-1 contributes to anti-tumoral activity of miR-101-3p.

## Background

Colorectal cancer (CRC) is still the third most common cancer worldwide [1] despite recent advancements in screening and treatment. It is estimated that over 140,000 new cases of colon and rectal cancers were diagnosed in 2018 in the United States alone [2]. MicroRNAs (miRNAs) are small nucleic acids involved in the post-transcriptional regulation of gene expression, and have been implicated in the pathogenesis and prognosis of CRC [3–5].

miRNAs are usually encoded in the human genome as clusters. After nuclear processing, these molecules are exported to the cytoplasm and loaded into RNA-induced silencing complexes (RISC), directing them against binding sites in the 3’-UTR region of target mRNAs, based on the degree of complementarity. While a perfect match leads to mRNA cleavage, miRNAs with partial complementarity lead to translation blockade and/or mRNA degradation through multiple mechanisms [6]. In either case, miRNAs predominantly act to decrease target mRNA levels [7]. Since a miRNA:mRNA perfect match is not required for miRNA silencing of its targets, one miRNA can affect the expression of hundreds of target transcripts. Hence, deregulation of a single miRNA can lead to global alterations in gene expression in a given cell [8]. Aberrant expression of miRNAs contributes to tumorigenesis mainly by two mechanisms: repression of tumor suppressor genes or loss of repression of oncogenes [9]. In the first case, miRNAs become overexpressed and downregulate the expression of tumor suppressor genes; in the latter case, miRNAs become downregulated themselves while their oncogene targets are overexpressed due to reduced post-transcriptional silencing. This abnormal miRNA profile can facilitate proliferation and survival of tumor cells in malignancies such as CRC [10].

miRNAs controlling pluripotency of embryonic stem cells have been associated with tumorigenesis in diverse cancers, including CRC [11–13]. In fact, cancer cells and pluripotent stem cells share the ability to proliferate rapidly and virtually indefinitely [14]. Strikingly, reprograming of somatic cells into induced pluripotent stem cells share many similarities with the process of malignant transformation [15]. Therefore, miRNAs controlling stemness and differentiation of stem cells have potential to be used as targets for the study of uncontrolled proliferation in cancer. However, this has not yet been tested in the context of CRC, and functional data on the effects of these miRNAs in the survival of CRC cells is still lacking.

We hypothesized that miRNAs involved in the control of pluripotency and differentiation of stem cells can alter the proliferation and survival of CRC cells. With that in mind, we have selected a panel of 31 miRNAs that have their expression modulated during the differentiation of embryonic stem cells [16]. We then set out to identify the effects of these miRNAs on the proliferation and cell death in a human cellular model of CRC. Importantly, most of the miRNAs in this panel have been described to be differentially expressed in CRC (Table 1). Here, we identified three miRNAs that suppressed proliferation of CRC cells while also inducing significant cell death. Microarray analysis of miR-101-3p targets revealed modulation of relevant cancer-related pathways. We also provide further evidence that loss of miR-101-3p expression in colorectal cancer can confer proliferative advantage to malignant cells.

**Table 1:**
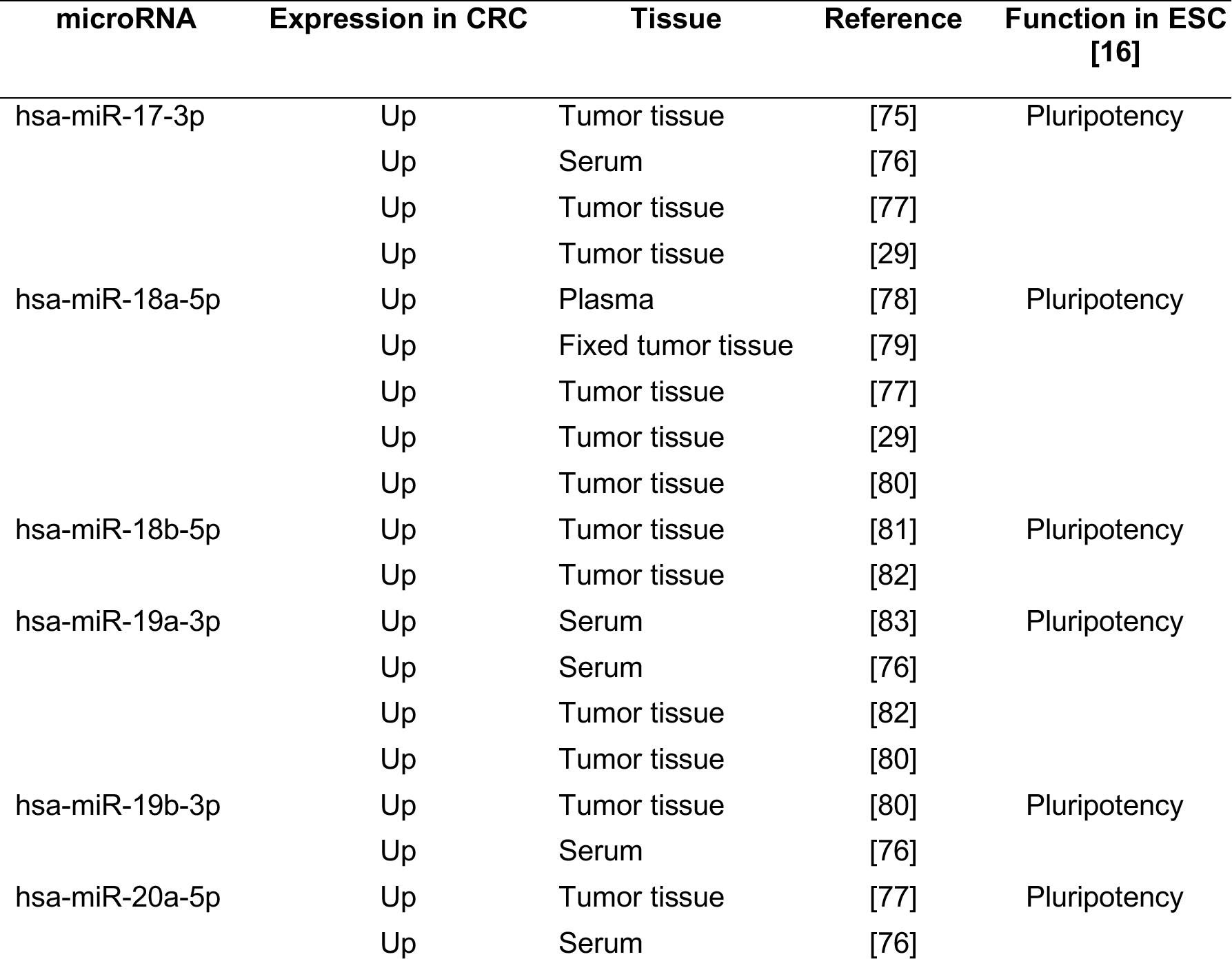
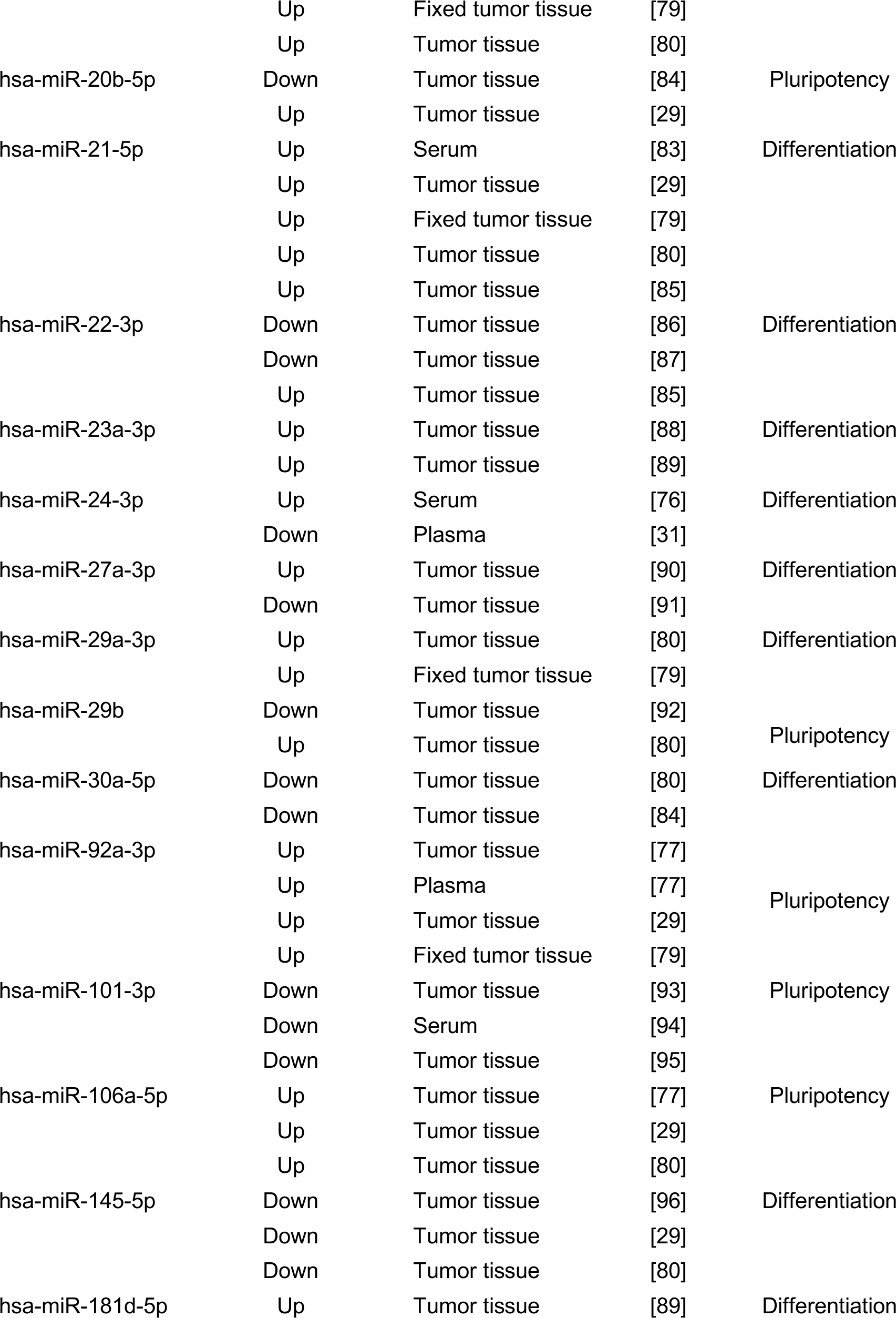
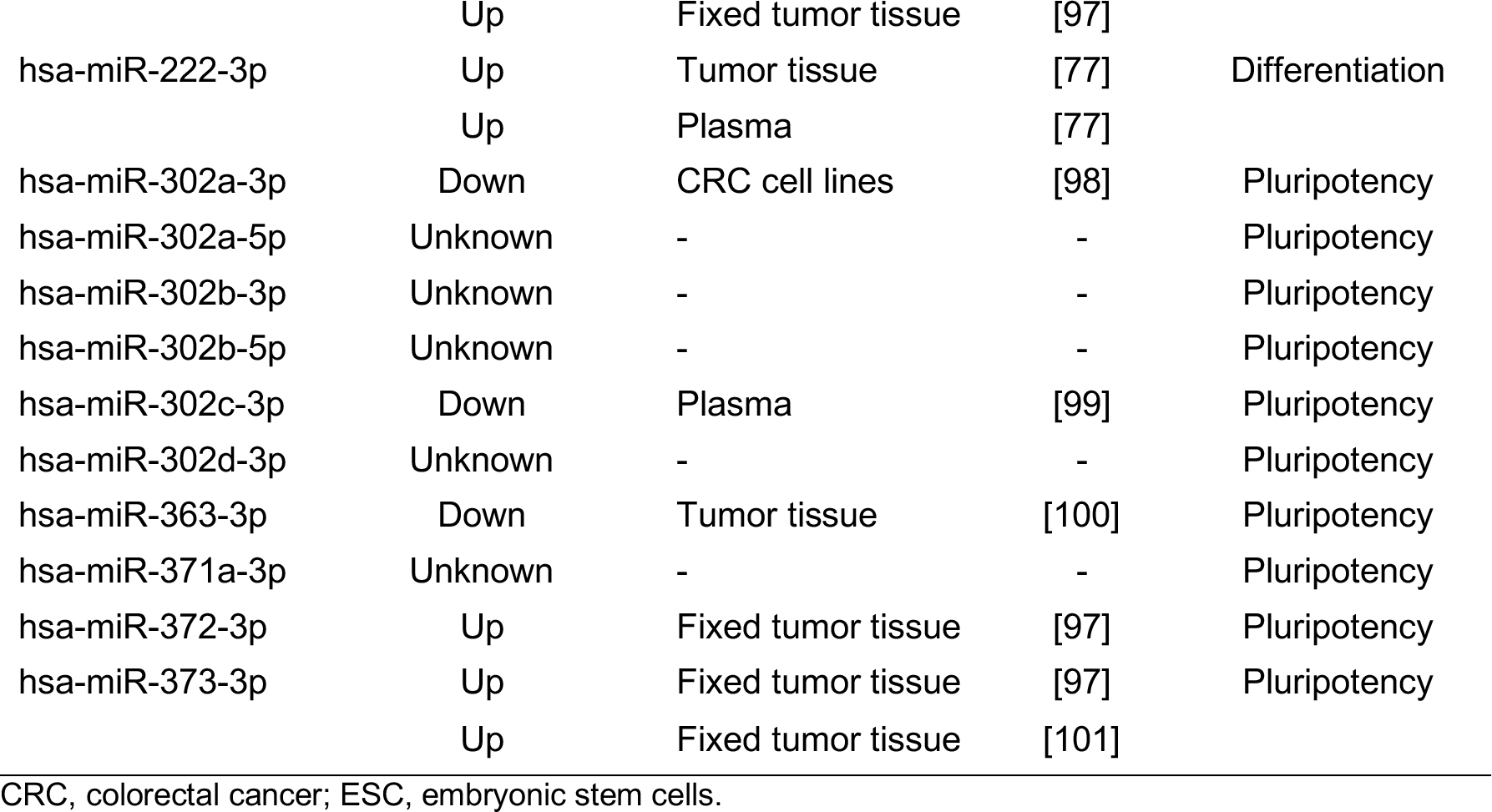
miRNAs differentially expressed in CRC and their function in embryonic stem cells (ESC).

| microRNA | Expression in CRC | Tissue | Reference | Function in ESC [16] |
| --- | --- | --- | --- | --- |
| hsa-miR-17-3p | Up | Tumor tissue | [75] | Pluripotency |
|  | Up | Serum | [76] |  |
|  | Up | Tumor tissue | [77] |  |
|  | Up | Tumor tissue | [29] |  |
| hsa-miR-18a-5p | Up | Plasma | [78] | Pluripotency |
|  | Up | Fixed tumor tissue | [79] |  |
|  | Up | Tumor tissue | [77] |  |
|  | Up | Tumor tissue | [29] |  |
|  | Up | Tumor tissue | [80] |  |
| hsa-miR-18b-5p | Up | Tumor tissue | [81] | Pluripotency |
|  | Up | Tumor tissue | [82] |  |
| hsa-miR-19a-3p | Up | Serum | [83] | Pluripotency |
|  | Up | Serum | [76] |  |
|  | Up | Tumor tissue | [82] |  |
|  | Up | Tumor tissue | [80] |  |
| hsa-miR-19b-3p | Up | Tumor tissue | [80] | Pluripotency |
|  | Up | Serum | [76] |  |
| hsa-miR-20a-5p | Up | Tumor tissue | [77] | Pluripotency |
|  | Up | Serum | [76] |  |
|  | Up | Fixed tumor tissue | [79] |  |
|  | Up | Tumor tissue | [80] |  |
| hsa-miR-20b-5p | Down | Tumor tissue | [84] | Pluripotency |
|  | Up | Tumor tissue | [29] |  |
| hsa-miR-21-5p | Up | Serum | [83] | Differentiation |
|  | Up | Tumor tissue | [29] |  |
|  | Up | Fixed tumor tissue | [79] |  |
|  | Up | Tumor tissue | [80] |  |
|  | Up | Tumor tissue | [85] |  |
| hsa-miR-22-3p | Down | Tumor tissue | [86] | Differentiation |
|  | Down | Tumor tissue | [87] |  |
|  | Up | Tumor tissue | [85] |  |
| hsa-miR-23a-3p | Up | Tumor tissue | [88] | Differentiation |
|  | Up | Tumor tissue | [89] |  |
| hsa-miR-24-3p | Up | Serum | [76] | Differentiation |
|  | Down | Plasma | [31] |  |
| hsa-miR-27a-3p | Up | Tumor tissue | [90] | Differentiation |
|  | Down | Tumor tissue | [91] |  |
| hsa-miR-29a-3p | Up | Tumor tissue | [80] | Differentiation |
|  | Up | Fixed tumor tissue | [79] |  |
| hsa-miR-29b | Down | Tumor tissue | [92] |  |
|  | Up | Tumor tissue | [80] | Pluripotency |
| hsa-miR-30a-5p | Down | Tumor tissue | [80] | Differentiation |
|  | Down | Tumor tissue | [84] |  |
| hsa-miR-92a-3p | Up | Tumor tissue | [77] |  |
|  | Up | Plasma | [77] | Pluripotency |
|  | Up | Tumor tissue | [29] |  |
|  | Up | Fixed tumor tissue | [79] |  |
| hsa-miR-101-3p | Down | Tumor tissue | [93] | Pluripotency |
|  | Down | Serum | [94] |  |
|  | Down | Tumor tissue | [95] |  |
| hsa-miR-106a-5p | Up | Tumor tissue | [77] | Pluripotency |
|  | Up | Tumor tissue | [29] |  |
|  | Up | Tumor tissue | [80] |  |
| hsa-miR-145-5p | Down | Tumor tissue | [96] | Differentiation |
|  | Down | Tumor tissue | [29] |  |
|  | Down | Tumor tissue | [80] |  |
| hsa-miR-181d-5p | Up | Tumor tissue | [89] | Differentiation |
|  | Up | Fixed tumor tissue | [97] |  |
| hsa-miR-222-3p | Up | Tumor tissue | [77] | Differentiation |
|  | Up | Plasma | [77] |  |
| hsa-miR-302a-3p | Down | CRC cell lines | [98] | Pluripotency |
| hsa-miR-302a-5p | Unknown | - | - | Pluripotency |
| hsa-miR-302b-3p | Unknown | - | - | Pluripotency |
| hsa-miR-302b-5p | Unknown | - | - | Pluripotency |
| hsa-miR-302c-3p | Down | Plasma | [99] | Pluripotency |
| hsa-miR-302d-3p | Unknown | - | - | Pluripotency |
| hsa-miR-363-3p | Down | Tumor tissue | [100] | Pluripotency |
| hsa-miR-371a-3p | Unknown | - | - | Pluripotency |
| hsa-miR-372-3p | Up | Fixed tumor tissue | [97] | Pluripotency |
| hsa-miR-373-3p | Up | Fixed tumor tissue | [97] | Pluripotency |
|  | Up | Fixed tumor tissue | [101] |  |

CRC, colorectal cancer; ESC, embryonic stem cells.

## Methods

### Cell culture and miRNA transfection

Human CRC cell line HCT116 (ATCC® CCL-247™) was cultivated using DMEM high-glucose supplemented with 10% FBS. Medium was changed every two days and cells were passaged by enzymatic treatment with TrypLE (ThermoFisher, Cat. No. 12604021) when 90-100% confluent. Cells were subcultured at 1:6 ratio into new flasks.

Synthetic miRNA mimics (pre-miRs) and an unspecific control (pre-miR control) were individually transfected into HCT116 cells by reverse transfection (**Supplementary Table 1**). In summary, 50uL of culture medium containing 8×10^3^ cells was added to wells of 96-well plates pre-filled with a mixture composed of 0.15 uL Lipofectamine RNAiMax (ThermoFisher, Cat. No. 13778150) and miRNAs in 50uL serum-free culture medium (50nM final miRNA concentration). Medium was changed 24h post-transfection, and cells were kept in culture for 4 additional days for proliferation assay. For gene expression analysis, 8×10^4^ cells were seeded in 6-well plates 18-24h before miRNA transfection. Transfection protocol was adjusted for a final volume of 1 mL. Cells were collected 72 h post-transfection for RNA extraction, used for qPCR and microarray analyses.

### Proliferation/Cell Death Assay

For proliferation assay, medium was removed after 4 days in culture and replaced by a 1.25 ug/mL solution of membrane-impermeant Propidium Iodide (PI) and 1uM of the membrane-permeant Hoechst 33342 (Hoechst) DNA stains, in final volume of 100 µL PBS. After an incubation period of 10 min, images were acquired by automated fluorescence microscopy using a High Content Screening platform (ImageXpress; Molecular Devices Inc.), under 10X objective. Excitation and emission channels used were 377/447 nm and 531/593 nm for PI and Hoechst, respectively. Nuclei of live cells (i.e. with intact membranes) were stained only by Hoechst, whereas nuclei of dead cells were stained by PI as well. Acquisition and analysis of images were performed using the platform’s built-in software MetaXpress, using the Live/Dead Application Module. For each well of a 96-well plate, nine fields were acquired and all cells within this area were quantified. ANOVA test with Dunnett post-test was used to detect differences between cells in wells transfected with control and miRNA mimics, in terms of total number of Hoechst-positive cells (proliferation) and percentage of PI-positive cells (death).

### RNA extraction and RT-qPCR

Total RNA was extracted from cells 72 h post-transfection using Trizol reagent (ThermoFisher, Cat. No. 15596018), following manufacturer’s instructions. cDNA was generated by reverse transcription using 1 ug of RNA as starting material, using the High Capacity cDNA Reverse Transcription kit (ThermoFisher, Cat. No. 4368814), following manufacturer’s instructions. Real-time qPCR reactions were performed using SYBR Green PCR master mix (ThermoFisher, Cat. No. 4309155) and in-house primers **(Supplementary Table 2**) using 10ng of cDNA. Relative gene expression was calculated using the 2^−∆∆CT^ method. *GAPDH* was the normalizer housekeeping gene and Control was used as reference sample. All experiments were performed in 3 biological replicates. t-test was used to detect differences between treatments and control.

### Oligonucleotide Microarray and Bioinformatics Analyses

Whole Human Genome Microarray Kit 4×44K (Agilent, Cat. No. G4112F) was used to detect mRNA expression levels in cells transfected with control and miR-101-3p transfected HCT116 cells, following manufacturer’s instructions. Differential expression of 41,000+ unique transcripts was analyzed using bioinformatics package Bioconductor (Gentleman, Carey et al. 2004). Results were normalized using LIMMA package [17]. Transcripts were considered differentially expressed when fold change was higher than 0.5 and p<0.05, using moderate T test. False Discovery Rate (FDR) test was used to adjust p values.

Predicted targets of miR-101-3p were obtained from the TargetScan database [18]. In order to carry a pathway enrichment analysis, we used the whole set of predicted targets that were also downregulated by miR-101-3p in our microarray analysis. This set of transcripts were analyzed using the Database for Annotation, Visualization and Integrated Discovery (DAVID) [19], restricting the analysis to pathway data from the Kyoto Encyclopedia of Genes and Genomes (KEGG) [20].

## Results

### microRNAs induce or halt proliferation of colorectal cancer cells

Several miRNAs have been reported to be differentially expressed in CRC tissue when compared to normal adjacent tissues, or between the serum of CRC patients and healthy controls. However, discrepant results are often found by different authors for several microRNAs (Table 1). Additionally, the functional outcomes of up- and downregulation of specific miRNAs in colorectal cancer cells remain to be fully evaluated.

To investigate the effects of miRNAs on the proliferation and survival of CRC cells, we performed a focused screen in HCT116 cell line. Cells were transfected with 31 synthetic miRNA duplexes mimics and cultured for 4 days. These so called pre-miR molecules are small, double-stranded RNA molecules designed to mimic endogenous mature miRNAs. Chemical modifications induce loading of the correct strand into RISC (**Supplementary Table 1**). Upon delivery via lipofection, one strand of the pre-miR molecule is loaded into RISC complexes, where it can modulate expression of target mRNAs, mimicking the effects of native miRNAs. Total number of live and dead cells was determined by fluorescence staining of transfected cells and imaging using a High content screening (HCS) platform. Image analysis of transfected cells allowed us to simultaneously identify miRNAs affecting cell proliferation and/or death of CRC cells (Figure 1; **Additional File 1**).

**Figure 1:**
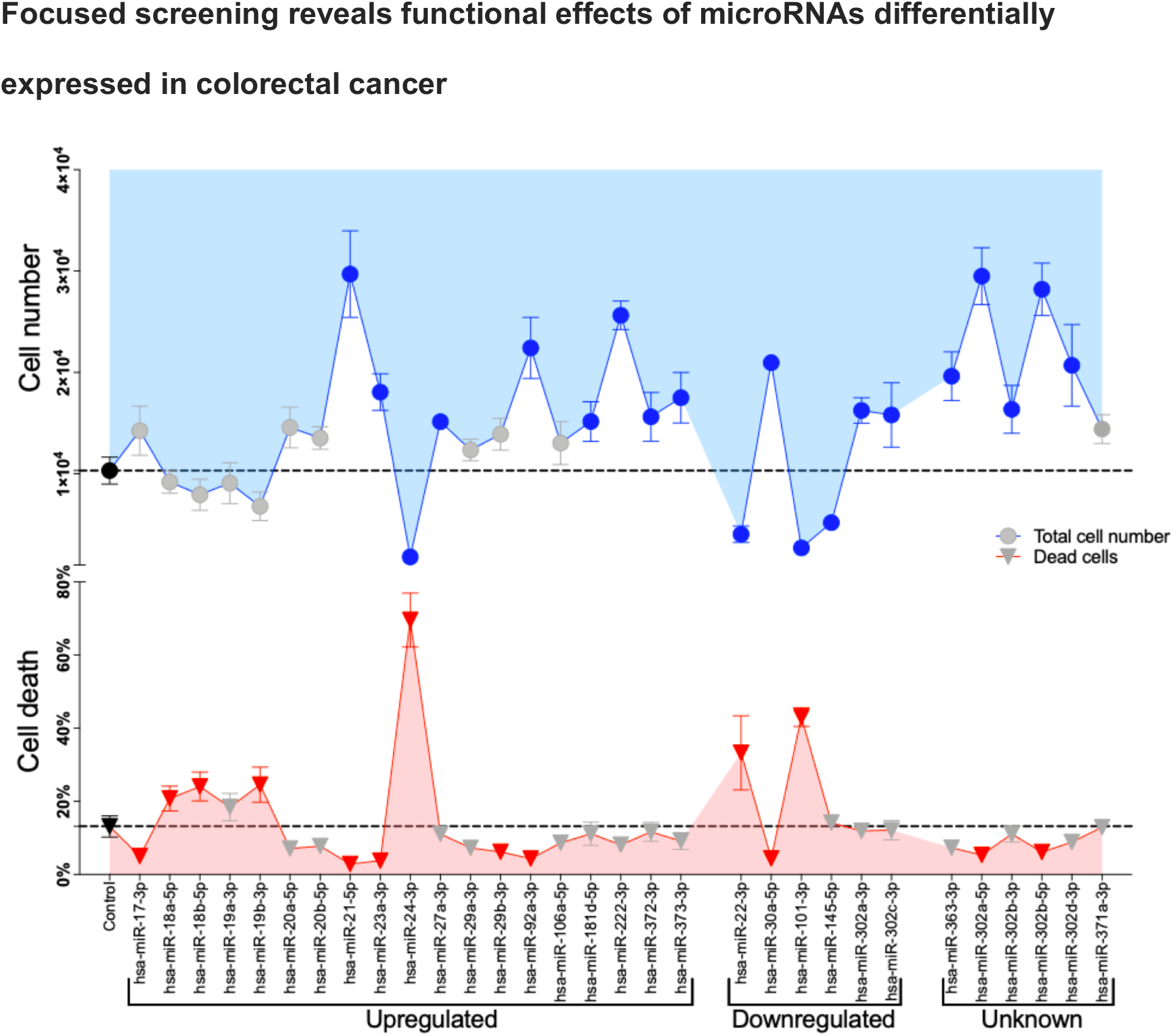
miRNAs differentially expressed in CRC modulate the proliferation of HCT116 cells. HCT116 were treated with 50nM of miRNA mimics for 4 days in 96-well plates. Total number of cells and number of dead cells were quantified by high-content screening analysis. Data is expressed as mean ± SD. Gray symbols indicate no significant differences.

Four days following transfection of miRNAs mimics on HCT116 cells, 16 of the miRNAs induced proliferation significantly (i.e. higher total cell counts, as compared to control), while only 4 repressed it. On the other hand, 8 miRNAs reduced cell death (i.e. lower percentage of dead cells, as compared to control), while 6 induced it. Eight miRNAs described as upregulated in CRC tissues or serum samples induced significant increase in cell proliferation (miR-21-5p, -23a-3p, -27a-3p, -92a-3p, -181d-5p, -222-3p, -372-3p, and -373-3p), whereas only one CRC-upregulated miRNA inhibited proliferation (miR-24-3p). Cancer and pluripotency related miRNAs belonging to clusters miR-106a~363, miR-17~92 and miR-302 induced significant increase in proliferation. Notably, mimics of miR-22-3p, miR-24-3p, and miR-101-3p, all described as CRC-downregulated miRNAs, simultaneously reduced cell proliferation and induced cell death, significantly, when compared to control. We decided to focus on miR-101-3p for further experiments due to described involvement of this miRNA in CRC.

### miR-101 downregulates signaling pathways controlling cell survival

Transfection of miR-101-3p in HCT116 cells led to reduction of cell numbers and increased cell death (Figure 2a). To identify potential post-transcriptional regulatory mechanisms mediating the observed functional effects of miR-101-3p, we performed a gene expression analysis using oligonucleotide microarray of HCT116 transfected with miR-101-3p mimics. A total of 4,826 transcripts were significantly downregulated by miR-101-3p (Figure 2b). In silico predictions of miR-101-3p targets from the TargetScan database [18] were crossed with our experimental data to identify transcripts most likely to be directly regulated by miR-101-3p (Figure 2c). Twenty percent (198 out of 947) of miR-101-3p predicted targets were downregulated in HCT116 treated with miR-101-3p. Moreover, 47 of these targets had also been experimentally validated by diverse studies cataloged by miRTarBase [21] (Figure 2d), featuring Wnt and apoptosis-related genes. This subset of predicted target transcripts downregulated upon introduction of miR-101-3p in HCT116 CRC cells, and previously experimentally validated in independent studies, represent high-confidence targets **(Additional File 2)**.

**Figure 2:**
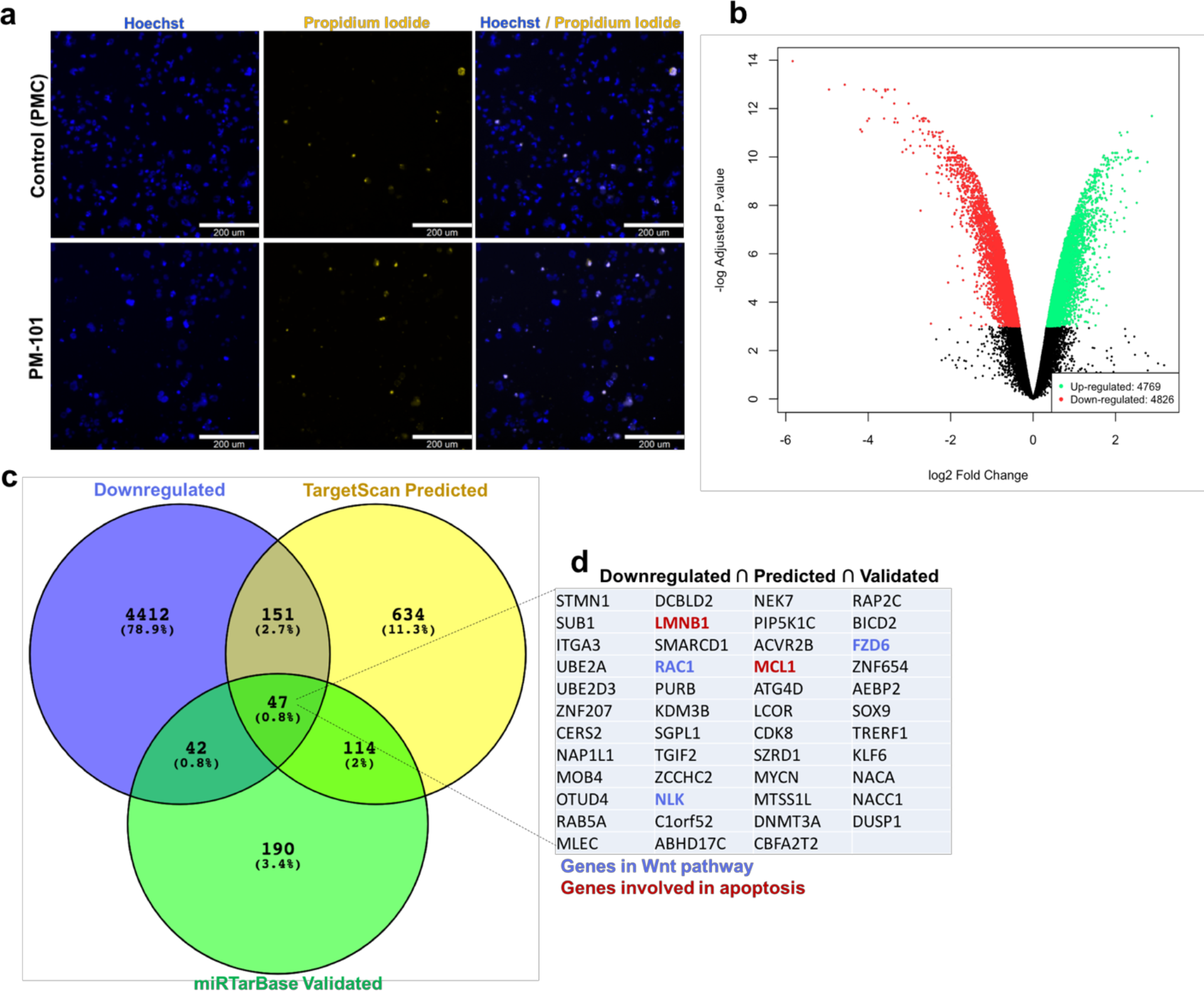
Differential gene expression in HCT116 treated with miR-101-3p. a) Representative fields of HCT116 treated with miR-101-3p for 4 days, showing increase in cell death and decrease of overall proliferation; b) Volcano plot showing downregulated (red) and upregulated (green) transcripts as a function of fold-change and P-value after 3 days of treatment; c) Experimentally downregulated targets are crossed with predicted and validated targets to identify high confidence targets; d) Functionally validated targets (miRTarBase) that were downregulated by miR-101-3p in microarray analysis.

In order to extract information regarding cellular processes that could be post-transcriptionally modulated by miR-101-3p, we performed a pathway enrichment analysis using Database for Annotation, Visualization and Integrated Discovery (DAVID) [19] by entering the set of predicted miR-101-3p targets that were downregulated in our microarray (198 transcripts). Perhaps not surprisingly, “Pathways in Cancer” and “Wnt” had the highest enrichment of targets associated with miR-101-3p expression. Kyoto Encyclopedia of Genes and Genomes (KEGG) (Kanehisa and Goto 2000, Kanehisa, Furumichi et al. 2017) pathway data was used to organize genes lists according to function. (Table 2).

**Table 2:**
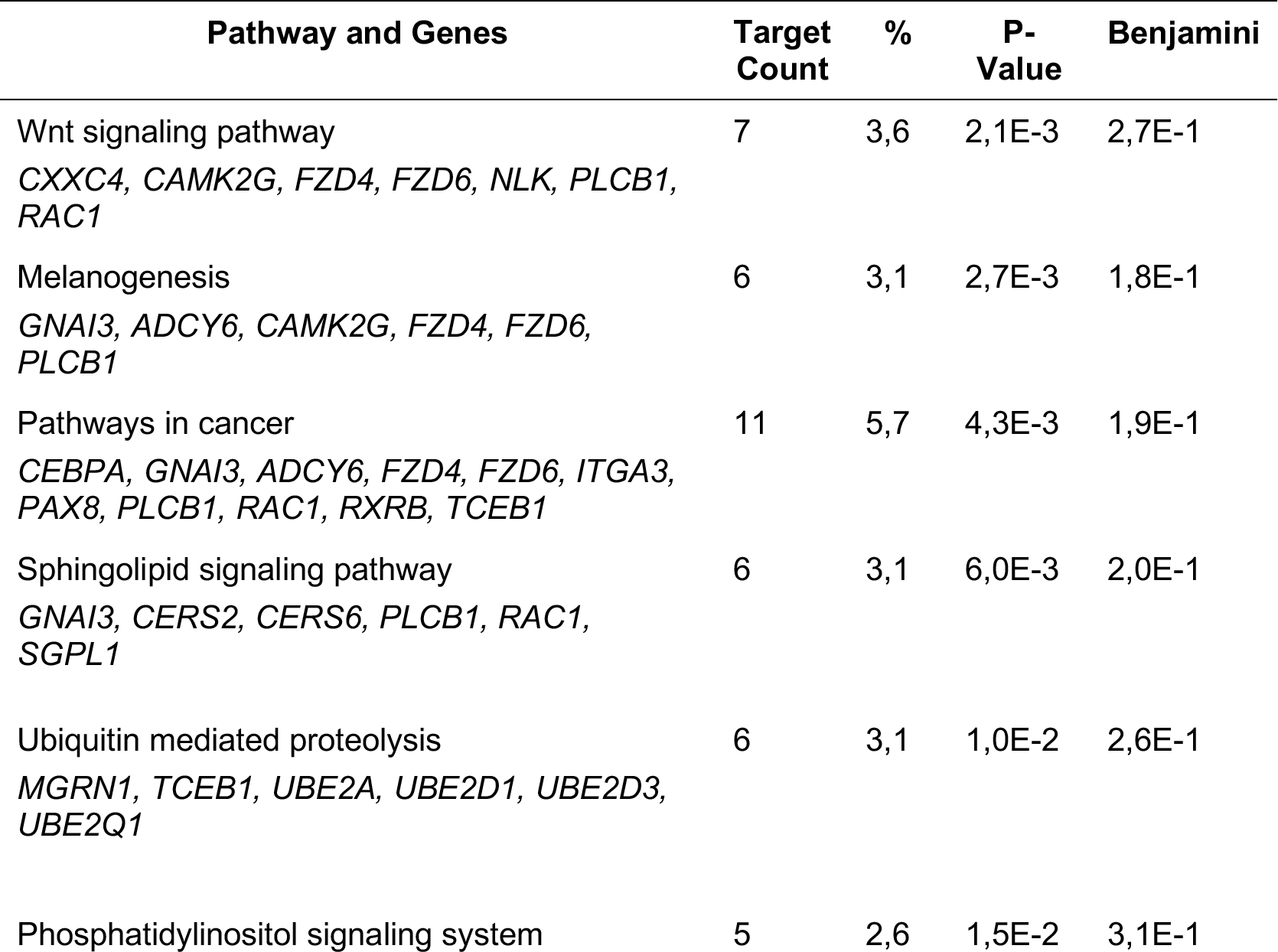
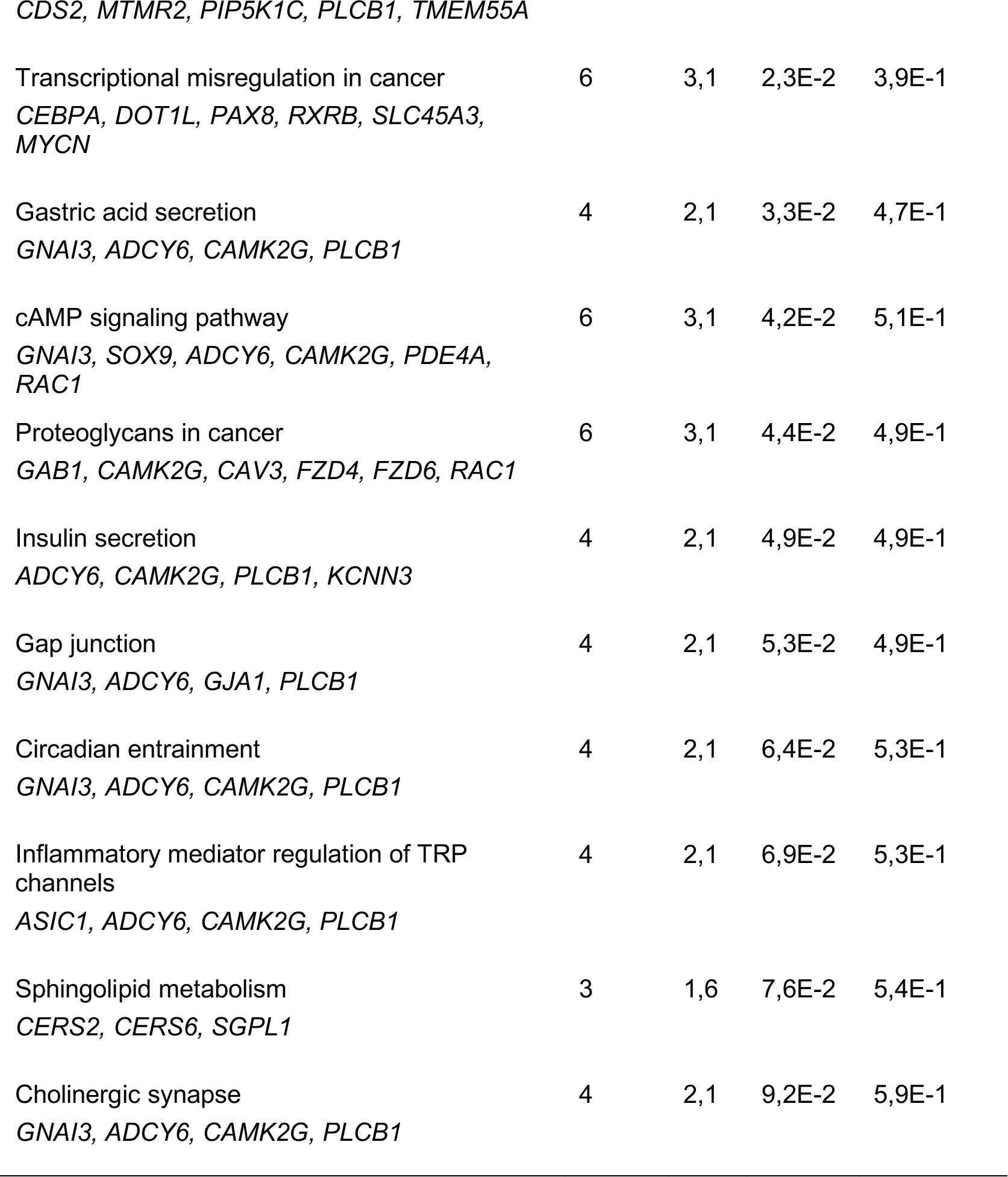
KEGG signaling pathways modulated by predicted miR-101-3p targets downregulated experimentally in HCT116 cells.

Next, we analyzed the expression of several putative miR-101-3p direct and indirect targets in HCT116 cells (Figure 3a). First, we analyzed the expression of several oncogenes and cell cycle related transcripts. Levels of tumor suppressors Phosphatase and Tensin Homolog (*PTEN*) and Cyclin Dependent Kinase Inhibitor 1C (*CDKN1C*) did not differ significantly from control treated cells. However, miR-101-3p expression inhibited v-myc myelocytomatosis viral oncogene homolog (*MYC*) mRNA, which can help explain, at least partially, the observed halt in cell proliferation of transfected HCT116 cells. Similarly, we observed repression of Enhancer of Zeste 2 Polycomb Repressive Complex 2 Subunit (*EZH2*), which is a predicted target of miR-101-3p that has been linked to oncogenesis in CRC [22, 23]. Analysis of canonical Wnt pathway genes reveled upregulation of ß-catenin mRNA (*CTNNB1*) whereas Glycogen Synthase Kinase 3 Beta (*GSK3B*) and Adenomatous Polyposis Coli (*APC*) remained unaltered, in spite of both *GSK3B* and *APC* being predicted targets of miR-101-3p.

**Figure 3:**
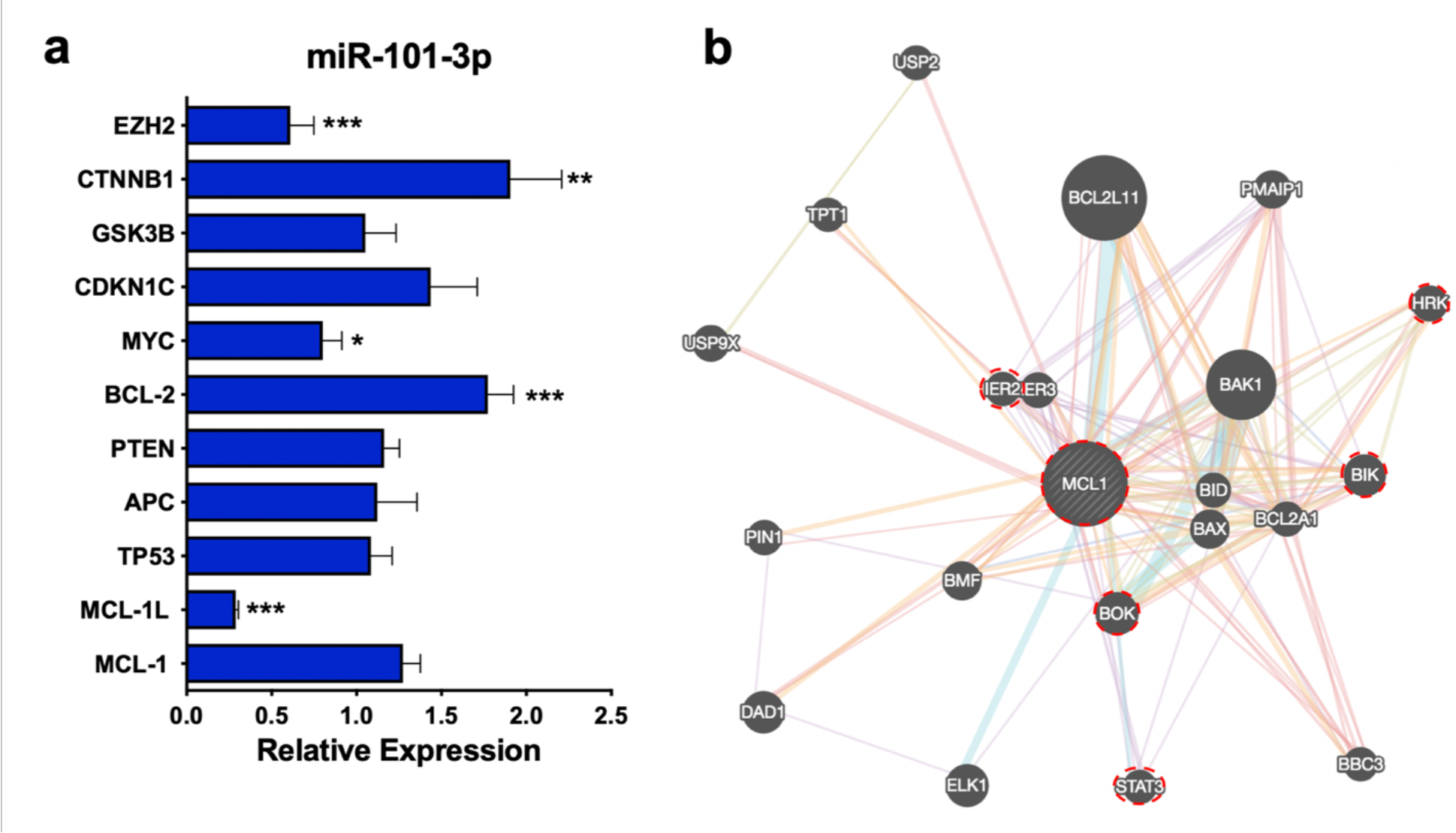
Genes downregulated by miR-101-3p in HCT116. a) Expression of potential direct and indirect targets of miR-101-3p in HCT116 cells relative to control; b) MCL-1 protein interaction network. Red circles indicate genes downregulated by miR-101-3p in microarray data. Data is expressed as mean ± SD. N=3. * p<0.01, **p<0.001, *** p<0.0001

We found that Myeloid cell leukemia-1 (*MCL1*), a member of the BCL-2 family of tumor suppressors and a predicted target of miR-101-3p, was downregulated in our microarray data. *MCL1* gene encoded three isoforms: one long, anti-apoptotic MCL-1L, and two shorter pro-apoptotic MCL-1S and MCL-1ES [24]. Downregulation of *MCL1* mRNA observed in the microarray was confirmed by qPCR. Interestingly, only the anti-apoptotic isoform of MCL-1 was downregulated by miR-101-3p **(MCL-1L**, Figure 3a). Additional apoptosis-related genes that closely interact with MCL-1 were also downregulated by miR-101-3p in our microarray data (Figure 3b).

## Discussion

In the present study, we evaluated the effects of 31 miRNAs on the proliferation and survival of a colorectal cancer cell line. Twenty miRNA mimics significantly altered HCT116 total cell numbers compared to control. Mimics of miR-22-3p, miR-24-3p, and miR-101-3p significantly reduced cell proliferation whilst inducing significant cell death when compared to control.

Differential expression of miRNAs is a common feature of many malignancies. Upregulation of oncomiRs and, conversely, downregulation of tumor suppressor miRNAs is believed to play a role in the proliferation and survival of cancer cells [9, 25]. Perhaps not surprisingly, around 30-50% of miRNAs are located at instable, cancer-associated genomic regions, and fragile sites [26, 27], which contributes to their aberrant expression profiles. miRNAs controlling pluripotency and differentiation of stem cells have been shown to be involved in tumorigenic processes and cancer stem cell derivation [28].

The panel of miRNAs tested in the present study represent miRNAs downregulated or upregulated during the differentiation of embryonic stem cells [16]. It is hypothesized that during the differentiation process downregulated miRNAs are involved in the maintenance of stemness properties, whereas miRNAs upregulated are involved in the induction of differentiation. More importantly, the miRNAs tested also represent the most frequently reported as differentially expressed miRNAs in CRC (Table 1).

Zhu and colleagues identified 38 miRNAs differentially expressed in tumor tissues of CRC patients [29]. Among the 30 miRNAs found upregulated in that study, miR-21-5p, miR-20b-5p, miR-106a-5p, miR-92a-3p and miR-17-3p were also included in our study and, except for miR-18b, they all stimulated cell proliferation of HCT116 cells in our screening, albeit only significantly for miR-21-5p and miR-92-3p. On the other hand, miRNAs that significantly reduced cell proliferation and induced cell death in our study had previously been described as poorly expressed in CRC samples: miR-22-3p [30], miR-24-3p [31], and miR-101-3p [32]. Ng *et al.* also detected differential expression of several miRNAs tested in our screening [33]. Among the concordantly upregulated miRNAs in serum and CRC biopsies in that study, miR-92-3p and miR-222-3p were also tested in our screening and induced significant increase in proliferation of HCT116 cells.

miRNAs can be expressed from polycistronic clusters wherein several miRNAs stem from the same primary transcript [34]. In this study, we investigated miRNAs belonging to cluster miR-17~92 (miR-17-3p, -18a-5p, -19a-3p, -19b-3p, -20a-5p, -92a-3p), cluster miR-106a~363 (miR-18b-5p, -20b-5p, -106a-5p, -363-3p) and cluster miR-302 (miR-302a-3p, -302a-5p, -302b-3p, -302b-5p, -302c-3p, -302d-3p). These clusters are abundantly expressed in pluripotent stem cells and are involved in stemness maintenance [35] while also being associated with deregulated proliferation and malignancies [36, 37]. Overall, the cluster miR-17~92 induced proliferation in our screening. This cluster is located at chromosome 13q31, one of the regions associated with CRC progression. Previous work has demonstrated that gain of the region containing this cluster leads to increased expression of the corresponding miRNAs in CRC tumor samples [37]. Taken together, the proliferation profile observed in our study points to a proliferative advantage for augmented expression of cluster miR-17~92 in CRC.

The pluripotency-associated cluster miR-302 induced marked proliferation of HCT116 cells in our study. Overexpression of this cluster is sufficient to reprogram somatic cells to pluripotency [38]. However, it has been suggested that although these miRNAs activate a pluripotency program in the target cells, they do so while also protecting cells from malignant transformation [39]. Previous work corroborating this idea has suggested that overexpression of miR-302 cluster actually can rescue malignant cells by reducing their proliferative profile and invasiveness [40]. A contrasting study indicates that overexpression of miR-302 cluster in cancer cells actually leads to a more invasive and undifferentiated cancer state [28]. Although our data supports the latter hypothesis, more studies in CRC models will be needed to address the context-dependent functions of miR-302 cluster in this malignancy.

miR-101-3p is one of the miRNAs downregulated during the differentiation of embryonic stem cells [16]. It markedly reduced cell proliferation and promoted cell death in our screening. Similar to our results, Chen and colleagues also demonstrated that overexpression of miR-101-3p in CRC in vitro models (HT-29 and RKO colon cancer cell lines) reduced proliferation and viability and simultaneously sensitized cells to 5-FU inhibition [41]. In fact, re-expression of miR-101-3p has been associated to in vitro sensitization of CRC cells to chemotherapy where miR-101-3p overexpression led to enhanced activity of paclitaxel and doxorubicin in HT-29 cells [42].

miR-101-3p has been found to act as a tumor suppressor in several malignancies, such as liver [43], glioblastoma [44], breast [45], endometrial [46], and colorectal [47].

Downregulation of this miRNA is so frequently found in solid tumors that some authors propose to use miR-101-3p expression as prognostic biomarker and therapeutic target [48–51]. miR-101-3p expression is commonly found downregulated in comparison to healthy adjacent tissues and, in some instances, its expression can predict poor prognosis and overall survival in CRC [32, 41, 52].

Epigenetic factors play an important role in CRC pathogenesis and progression [53]. Here we have shown that miR-101-3p significantly repressed expression of EZH2, a member of the Polycomb Repressor Complex 2 (PCR2) which catalyzes methylation of lysine 27 of histone H3 (H3K27me3). This complex modifies the chromatin structure to favor a proliferative program by bypassing the Ink4a/Arf-pRb-p53 pathway [54]. EZH2 promotes proliferation of CRC cells, and its silencing by siRNA leads to reduced cancer cell survival [22]. Recent data has suggested the existence of a negative feedback loop between EZH2 and miR-101-3p. Treatment of CRC cells with an anti-cancer substance named methyl jasmonate led to apoptosis and inhibited expression of EZH2 while upregulated miR-101-3p expression [55]. Furthermore, EZH2 has been linked to epigenetic inactivation of WNT5A, a proposed tumor suppressor, during TGF-ß-induced epithelial-mesenchymal transition in an in vitro model of CRC [56]. EZH2 might also be involved in CRC chemotherapeutic efficacy. EZH2 repression increased the efficiency of EGFR inhibitors in vitro [23]. Similarly, Yamamoto and colleagues have shown that EZH2 expression was associated with survival in CRC patients undergoing anti-EGFR therapy [57].

Hypermethylation has been associated with CRC pathogenesis in several studies [58-60] [reviewed in [61]]. Aberrant hypermethylation phenotype of tumor suppressor genes by DNMT3a activity has been reversed by expression of miR-101-3p in a model of lung cancer, where DNMT3 repression led to promoter hypomethylation and re-expression of tumor suppressor CDH1 [62]. Perhaps not surprisingly, our microarray data revealed downregulation of both DNMT3a and DNMT3b in HCT116 cells treated with miR-101-3p mimics. Similarly, Toyota and colleagues demonstrated that miR-34b and miR-34c were epigenetically silenced in HCT116 cells, and its expression could be rescued by treatment with 5-aza-2’-deoxycytidine, a DNA demethylation agent. Moreover, they showed that CpG methylation of miR-34b/c was a common feature of different CRC lines [63]. Although authors did not investigate the methylation levels of miR-101-3p locus, CpG methylation represents a possible mechanism for repression of other miRNAs in CRC.

Our microarray data helps shed light on the involvement of Wnt pathway in CRC. Inhibition of Wnt pathway results in reduced proliferation in several cancers, including CRC [64]. We found B-catenin mRNA, the main mediator of canonical Wnt pathway, to be overexpressed in HCT116 cells in response to miR-101-3p mimics. However, overexpression of *CTNNB1* as measured by qPCR could reflect the mutated status of this gene in HCT116 cell line. Mouradov and colleagues performed an extensive whole-exome sequencing and SNP microarray analysis of 70 CRC cell lines, which revealed *CTNNB1* mutated status of several of them, including HCT116 [65]. It is interesting to speculate that miR-101-3p may interfere with the non-canonical Wnt pathway, given that genes downregulated in our microarray most likely reflect this hypothesis (*CXXC4, CAMK2G, FZD4, FZD6, NLK, PLCB1, RAC1*). For instance, expression of Nemo-like kinase (NLK) has been demonstrated to be necessary for cell cycle progression in CRC in vitro [66].

In addition to inhibiting cell proliferation, miR-101-3p also remarkably induced cell death in treated cells. Several pathways have been implicated in the induction of apoptosis by miR-101-3p in different cancer cell models [67–69]. Microarray analysis provided some clues on what genes can be modulated in order to warrant such effect on cell survival. MCL-1 is a member of the BCL-2 superfamily of apoptosis regulators, and it is one of the most frequently amplified genes in cancers [70]. MCL-1 gene encodes three isoforms: the long, anti-apoptotic MCL-1L, and two shorter pro-apoptotic MCL-1S and MCL-1ES [24]. MCL-1 amplification accounts for resistance to the BCL-2/BCL-xL inhibitor ABT-737 (Chen et al., 2007, Keuling et al., 2009, van Delft et al., 2006). MCL-1 associates with mitochondrial membrane-associated proteins, Bak and Bax, preventing them from heterodimerizing with apoptotic members of the BCL-2 family to promote apoptosis cascade [71]. More strikingly, a recent study has demonstrated that degradation of MCL-1 is necessary for effective therapeutics against CRC [72]. Similarly, inhibition of MCL-1 by miR-101-3p has been implicated in the apoptosis-inducing effect of anti-cancer drug doxorubicin in hepatocellular carcinoma [73]. A similar inhibitory mechanism between miR-101-3p and MCL-1 has been reported for endometrial cancer as well [46]. The specific inhibition of only the anti-apoptotic MCL-1 isoform in our study highlights a novel mechanism by which miR-101-3p can induce apoptosis and cell death in our screening, and points to a possible therapeutic target for oligonucleotide-based therapies.

Overall, the cell proliferation profile observed in our model of CRC points to an interesting tendency: miRNAs overexpressed in CRC augment cell proliferation and, conversely, miRNAs poorly expressed in CRC reduce cell proliferation and survival. Additionally, microRNAs characteristic of pluripotent stem cells tend to confer a proliferative advantage to CRC cells. This phenomenon suggests the existence of potential functional advantages of the differential expression of miRNAs observed in colorectal cancer. Since selective pressure within tumor tissue favors accumulation of genetic alterations that support survival [74], it is tempting to speculate that miRNAs consistently described as downregulated in CRC could have been selectively repressed due to their effects on proliferation, such as seen in our study.

## Conclusions

Taken together, the results provide additional evidence of functional outcomes resulting from differential expression of miRNAs in CRC. Additional studies will be necessary to elucidate the mechanisms by which miRNAs differentially expressed in CRC promote these effects on proliferation, and the present study points to interesting miRNAs to pursuit. Additionally, miR-101-3p appears to target multiple transcripts that act synergistically to promote cell death and halt proliferation of CRC cells in vitro, mainly by targeting Wnt pathway. More specifically, we provide novel evidence linking inhibition of MCL-1 by miR-101-3p as a potential mechanism for antitumoral activity of this miRNA.

## Additional Files

Additional File 1: miRNA screening data and statistical analyses (.xlsx)

Additional File 2: Microarray data and statistical analyses (.xlsx)

## List of abbreviations

5-FU: 5-fluouracil
BCL-2: B-cell lymphoma-2
CRC: colorectal cancer
DAVID: Database for Annotation, Visualization and Integrated Discovery
EGFR: epidermal growth factor receptor
EZH2: Enhancer of Zeste 2 Polycomb Repressive Complex 2 Subunit
KEGG: Kyoto Encyclopedia of Genes and Genomes
MCL-1: Myeloid cell leukemia-1
NLK: Nemo-like kinase
miRNA: microRNAs
miR: microRNA
RISC: RNA-Induced Silencing Complex

## Declarations

### Ethics approval and consent to participate

Not applicable

### Consent for publication

Not applicable

## Availability of data and material

The datasets used and/or analysed during the current study are available under Additional Files.

## Competing interests

Authors declare no conflicts of interest.

## Funding

This work was supported by the Brazilian National Council for Scientific and Technological Development, CNPq (Fellowship Process No. 142491/2011-0), and by the São Paulo Research Foundation, FAPESP/Center for Cell-Based Therapy, CTC (Grant No. 2013/08135-2).

## Authors’ contributions

DS designed and performed experiments, data analyses and wrote manuscript; JB performed microarray analyses; IMSL and JLSS performed experiments; DTC provided study materials; RAP conceptualized screening approaches and supervised experiments. All authors read and approved final manuscript.

## Acknowledgements

Authors thank Dr. Joao Farias Guerreiro, MS. Amelia Araujo, Elizabete Audino and Claudia Magnani for their support in the execution of this project. Authors also thank Joey Owen and Erika Paulson for revising the final manuscript.

